# BLeaching In-cell Single-molecule burstS (BLISS) reveals dynamic fraction of HP1α clusters in chromocenters of undifferentiated embryonic stem cells

**DOI:** 10.64898/2026.04.26.720855

**Authors:** Khalil Joron, Eden Mishne, Eran Meshorer, Eitan Lerner

## Abstract

Confocal fluorescence microscopy measurements of dense cellular regions of interest (ROIs) in cells using fluorescence lifetime imaging microscopy (FLIM) provide detailed pixelated images. Yet, those pixels report ensemble- and time-averaged biomolecular data, due to the diffraction limit and acquisition times that are slower than typical biomolecular dynamics. The ability to acquire data on one biomolecule at a time within a given ROI can help recover some of the underlying biomolecular subpopulations that are otherwise masked out. Here, we present a simple approach to achieving single-biomolecule photon bursts in live cells, dubbed BLeaching In-cell Single-molecule burstS (BLISS), which does not necessarily require super-resolution modalities. Using BLISS, we show that mCherry-tagged heterochromatin protein 1α (mCherry-HP1α) in chromocenters of undifferentiated mouse embryonic stem cells (ESCs), exhibit photon bursts with millisecond durations arising from dynamic clusters. Fluorescence lifetimes of these bursts are substantially lower than the pixel-wise averaged values observed in FLIM, attributed to higher density in HP1α clusters. These higher density clusters are observed primarily in undifferentiated ESCs, while two days post retinoic acid (RA)-induced differentiation (2d-RA), these bursts are rarely observed. Using BLISS, we also detect dynamic clusters of the chromocenter-associated protein, CENP-V, and the nucleolar protein, nucleolin; however, in both cases the fluorescence lifetimes of their clusters closely resemble the pixel-wise averaged values recovered from FLIM. In conclusion, we introduce BLISS as a method for revealing rare and dynamic subpopulation in dense ROIs that are otherwise hidden by averaging in diffraction-limited imaging techniques. Importantly, using BLISS we demonstrate that denser-than-average HP1α clusters are hyperdynamic.

## Introduction

Single-molecule approaches reveal the heterogeneity and dynamics of various biologically meaningful readouts, one molecule at a time^1^. This feature is achieved by reporting the distribution of these single-molecule readouts and whether a single group or multiple distinct sub-groups can describe their distributions. Yet, resolving single-molecule fluorescence-based readouts in cells has been achieved so far by (i) reintroducing ultra-low concentrations of dye-labeled biomolecules back to the cell via transfection, transformation, electroporation, or microinjection^2–4^, (ii) measuring specifically-tagged biomolecules with bright fluorophores that can undergo controlled or stochastic photoswitching^5–14^, (iii) by tagging with specific binder biomolecules (e.g., DNA or protein tags), to which fluorescent substrates can bind^15–24^, (iv) combinations of the above methods with different super-resolution (SR) fluorescence microscopy techniques^6,7,25–32^, or (v) simply by tagging biomolecules of which there are only few within the cell, and only if they are well-separated, either biologically (e.g., telomeres in the nucleus, binding to specific chromosomal loci in highly inclined illumination optical, HILO, microscopy mode^33^, receptors atop the plasma membrane in total internal reflection, TIRF, microscopy mode^34–37^), or in regions of sufficiently low concentration^3,21,23,24^. Collecting single-molecule fluorescence-based information using intrinsically-tagged biomolecules expressed at endogenous levels combined with fluorescence microscopy modalities that many labs have at their disposal, such as confocal microscopy, would be an obvious advantage. Importantly, none of the aforementioned non-super-resolution approaches are suitable for tracking individual biomolecules in highly concentrated ROIs within cells.

Unlike in-cell studies, and particularly in dense ROIs, the literature is rich in single-molecule fluorescence spectroscopy studies in the test tube, outside the cellular context. There are several acquisition modalities for obtaining single-molecule fluorescence information, which can generally be divided into two categories: camera-based long-lasting observations (seconds-minutes) of individual molecules and point detection of single molecules diffusing into a confocal spot. In the former, TIRF, HILO, or light-sheet fluorescence microscopy are preferred over widefield microscopy, due to the more localized 2D sample slice that is segmented and imaged. In point detection, however, one sets the focused confocal spot at a given position in the sample space and collects molecules crossing it occasionally.

Indeed, in solution, one can collect single-molecule bursts of fluorescence if the concentration of dye-labeled molecules is sufficiently low (i.e., tens of pM) and the detected photon rate (i.e., the brightness of the dye) is sufficiently high relative to the experimental background^1^. However, in cells, especially in dense ROIs, it is difficult to ensure such low concentrations of fluorescently-tagged molecules. Since these ROIs are highly concentrated at cellular regions that yield high fluorescence intensity, it would be beneficial to detect single-molecule fluorescence bursts to resolve their readout distributions. We propose a rather simple approach, BLISS, inspired by photobleaching experiments^38–41^. BLISS achieves this using laser-scanning confocal microscopy and fluorescent protein (FP)-tagged proteins in living cells, employing continuous photobleaching^42,43^: the same focused laser illumination is used to continuously photobleach a small ROI and to acquire single-biomolecule fluorescence (Fig. 1a). We harness the balance between photobleaching, which depends on time, laser intensity, and concentration of fluorescently-active molecules, and the diffusion rates of fluorescently-tagged biomolecular assemblies and clusters out of the bleached ROI. Therefore, fluorescently-tagged assemblies that are sufficiently bright and diffuse into the imaged ROI from outside may be observed as photon bursts if the background is sufficiently low. Using BLISS, we demonstrate the acquisition of single-molecule fluorescence lifetimes of endogenously expressed mCherry-tagged proteins including HP1α, CENP-V and nucleolin, in mouse embryonic stem cells (ESCs)^44^. BLISS uncovers a hyper-dynamic fraction of HP1α clusters mostly in undifferentiated ESCs.

**Figure 1.**
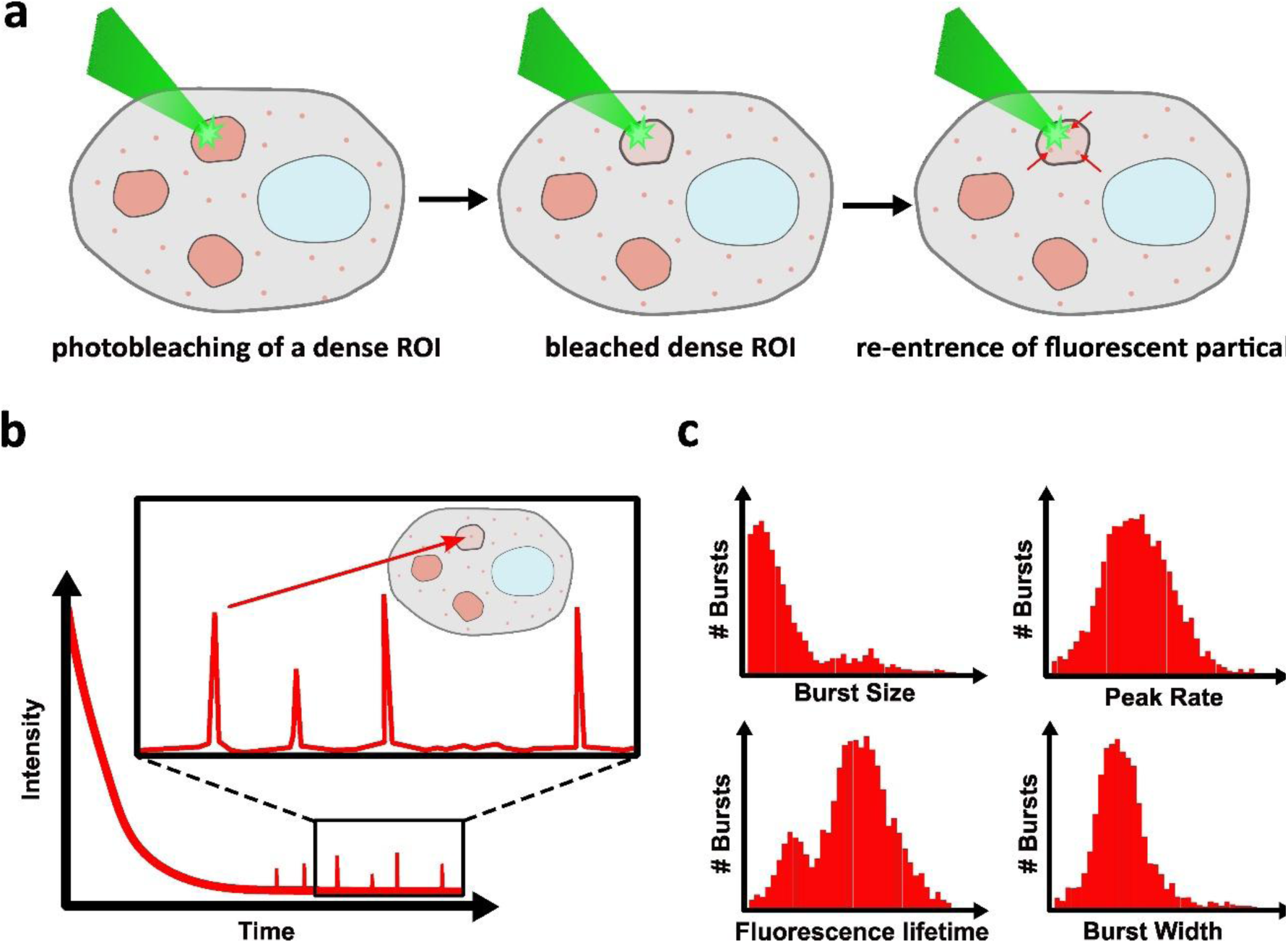
Illustration of BLeaching In-cell Single-molecule burstS (BLISS) principles. (a) Cells expressing a fluorescently-tagged protein that is part of a cellular ROI. Continuous photobleaching of the fluorescent tags in a given ROI, and re-entry of fluorescently-tagged proteins from outside the ROI into the photobleached site. (b) Time-trace of a measurement of fluorescence using a high-intensity focused laser, leading to both continuous photobleaching and the collection of fluorescence, including fluorescent bursts. The measurements start at high intensity, rapidly reaching steady-state background fluorescence, atop which photon bursts arising from diffusion of bright single particles are detected, through the laser focus and the photobleached ROI. (c) Mapping of burst-based readout histograms to identify different properties such as size, peak photon rate, width, and fluorescence lifetime.

### Main

Previously, we have shown that mCherry fluorescence lifetimes directly report elevated molecular densities, including those that occur in heterochromatin biocondensates or chromocenters, from mCherry-HP1α molecules^45^. Using FLIM we have also shown that pixel-wise densities change as ESCs differentiate and suggested that in undifferentiated ESCs, these biocondensates exhibit patches of different lifetimes, hence different densities^45^. However, what we were not able to report was whether pixel-wise lifetimes were by themselves averaging-out and masking underlying density heterogeneity. Based on recent work^45–49^, we suggest that the heterogeneity observed in FLIM arises from HP1α clusters interconnected through multiple co-interactors, such as major satellite repeat (MSR) RNA, linker H1 and other possible interactors acting as dynamic glue, including HP1 itself. Here, using BLISS in mouse ESC chromocenters, we recover the fluorescence lifetime distribution of individual HP1α clusters that re-enter the biocondensates during continuous photobleaching. We show that some HP1α clusters with millisecond translational dynamics exhibit lower fluorescence lifetimes than the pixel-wise averaged values recovered from FLIM, suggesting some HP1α clusters are denser than others. Remarkably, in early-differentiating cells following two-day RA (2d-RA) treatment, the amount of these HP1α clusters becomes significantly lower, implying that a highly mobile subpopulation of HP1α clusters is predominant in undifferentiated ESCs.

### BLISS approach and workflow

BLISS is achieved while knowing the background rate of the point detectors in use, the capability to quantify the actual fluorescence background arising from small, rapidly diffusing, high-concentration fluorescently-active molecules, as well as photon bursts with sufficiently high signal-to-background ratio (SBR). These photon bursts arise from brighter biomolecular clusters (Fig. 1b). In this approach, strong continuous photobleaching assists in reducing the concentration of the fluorescent molecule so that when a steady-state between photobleaching and molecular diffusion is achieved, the overall fluorescence signal becomes close to the detector background rate. At this point the signal is sufficiently low to identify bursts of photons arising from true molecular clusters above the background, better known as a single-particle fluorescence burst (Fig. 1b). Whether continuous photobleaching will lead to the formation of a dark hole at the imaged point strongly depends on the laser irradiance intensity, as well as on the diffusion rates of fluorescently-tagged biomolecules outside the photobleaching point, which provide fresh fluorescent molecules to be imaged. It is hence important to use single-photon point detectors with well-defined background rates, and a burst search and selection algorithm, such as the well-known consecutive photon sliding window approach^50^, which could define the actual background rate temporally, and abrupt photon bursts with photon rates that are sufficiently high relative to the background.

After acquiring a sufficient number of photon bursts for each chosen spot, a readout can be calculated for each burst, to yield the single-particle readout (Fig. 1c). Such readouts can be, for example, the number of photons per burst (i.e., burst size), the peak photon rate of the burst (i.e., equivalent to the molecular brightness), the duration of the burst (i.e., burst width), ratios of photon counts per burst detected on different detectors, or mean photon nanotimes, which report mean fluorescence lifetimes^51,52^. Such burst-wise quantities can be mapped to meaningful underlying insights, such as FRET efficiencies in FRET experiments^32,53^ or microenvironmental effects^54,55^, e.g., in FLIM^45^. Next, the distribution of these single-particle burst quantities for many recorded bursts can identify division into subgroups with distinct means.

### Simulating BLISS of rare clusters in the presence of diffusing molecules

In principle, the scenario presented in a BLISS experiment, after continuous photobleaching, leads to stable low fluorescence levels, which can be approximated by steady-state conditions. This scenario can be simulated in Brownian motion simulations of molecules diffusing through a confocal volume, simulated by a focused laser point spread function (PSF) profile, in which (i) there are many diffuse molecules with low molecular brightness values, and (ii) a few molecules with higher molecular brightness values, where their number can be compared to typical confocal-based single molecule fluorescence concentrations, a few tens of picomolars (pM).

From the simulation, a few key principles emerge for resolving bursts from rare bright species in the presence of abundant species. Detection sensitivity strongly depends on the laser intensity and the initial fluorophore concentration. Increasing the brightness contrast between the two populations markedly improves burst discrimination. Furthermore, burst detection is optimized by employing SBR instantaneous photon rate thresholds of at least 6 (minimal SBRs of 5; see Supplementary information for more details).

### Unmasking nanoscale HP1α heterogeneity in dense chromocenters of undifferentiated mouse ESCs using BLISS

Previously, we reported heterogeneity of fluorescence lifetimes of mCherry-HP1α in heterochromatin condensates, specifically chromocenters, of undifferentiated ESCs^45^. We also reported that in the 2d-RA condition, these condensates become more homogenous and less dense. The heterogeneity in fluorescence lifetimes and density was observed between pixels in FLIM measurements. HP1α is known to be found mainly in two states, (i) a form of HP1α loosely-bound to heterochromatin which exhibits a hyperdynamic fraction that is found especially in undifferentiated ESCs^38,56^, and (ii) a form bound to nucleosomes clusters of different densities^56^. It is suggested that segregation into multiple segments within a condensate, having different densities, has an important role in regulating processes inside heterochromatin biocondensates^57,58^. Nevertheless, it is still unclear what leads to the observed heterogeneity in heterochromatin densities, particularly when considering that FLIM reports mean fluorescence lifetimes of diffraction-limited pixels that may include a large number of underlying fluorescently-tagged molecules. Yet, if at least a fraction of the heterogeneity of fluorescence lifetimes found in FLIM arises from clusters of mCherry-HP1α with different densities we would expect to find more of these clusters in undifferentiated compared to 2d-RA ESCs, where the later shows high homogeneity of lifetimes in FLIM. Revealing the individual different contributors to the observed mean fluorescence lifetime per pixel can assist in explaining the observed heterogeneity. To address this, using BLISS we measure single clusters of mCherry-HP1α recurring inside puncta during its continuous photobleaching in both undifferentiated and 2d-RA ESCs (Fig. 2a-b). Then, using the consecutive photon sliding window burst detection approach, we analyze the fluorescence lifetime of photon bursts arising from these clusters.

**Figure 2.**
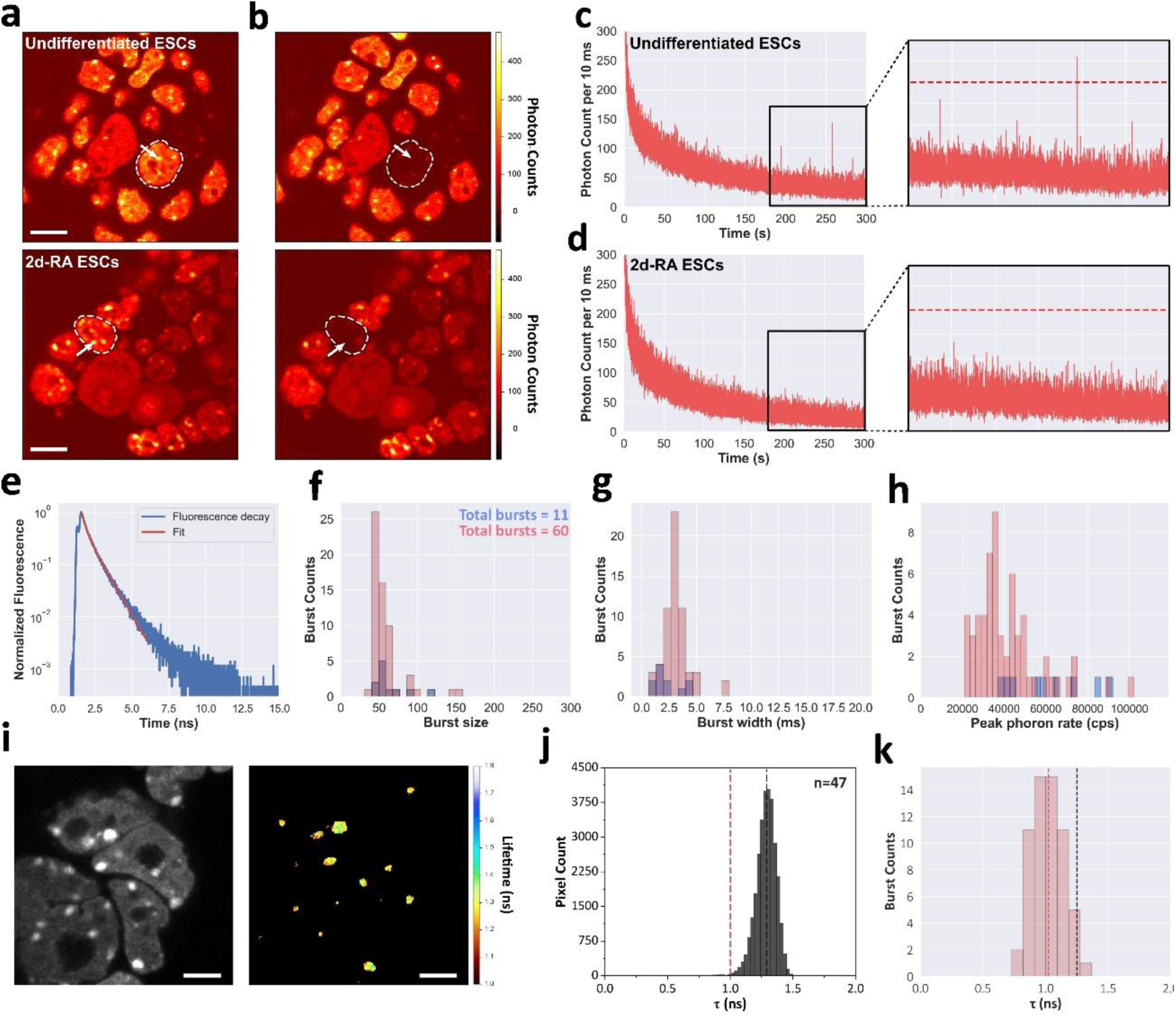
BLISS on mouse ESCs expressing mCherry-HP1α recovers clusters of HP1α with densities higher than the mean density recovered from FLIM. (a-b) Images of cell nuclei, focusing on a ROI (white arrows) in one nucleus before (a) and after (b) continuous photobleaching and collection of fluorescence, for both pluripotent (top) and early-differentiated (bottom) ESCs. (c-d) Examples of 300 s fluorescence time traces of mouse ESCs at *2i* stage (c) and at early differentiation stage - 2-days following differentiation onset (d) that exhibit continuous photobleaching, until reaching a steady-state with low counts serving as background levels, atop which photon bursts clearly appear in ESCs at *2i* and not at early differentiated states. A minimal single-to-background ratio (SBR) threshold of F=6 (indicated as red dashed line), is used for identifying these photon bursts and minimizing false detection of noise fluctuations about the mean fluorescence. (e) Readouts of fluorescence decay constructed from the photon nanotimes of all detected photons from the background, not including bursts (blue). Burst-wise histograms of (f) size, (g) width, and (h) peak photon rate. (i) Intensity image and FLIM of HP1ɑ puncta in nuclei of ESCs at *2i* stage. (j) Pixel-wise mean fluorescence lifetime histogram from FLIM of ESCs at the *2i* stage from puncta. (k) Burst-wise mean fluorescence lifetime histogram from BLISS of ESCs at the *2i* stage from puncta. Black dashed line – mean fluorescence lifetime recovered from FLIM. Red dashed line – mean fluorescence lifetime recovered from BLISS. Scalebars, 10 μm (a) and 5 μm (i).

Firstly, we retrieve steady-state background rates starting from 5.0 and ending at 1.0 kHz, for mCherry-HP1α measured in undifferentiated and 2d-RA ESCs during continuous photobleaching (Fig. 2c-d & Supplementary Fig. 1). We acquire the fluorescence decay from inter-burst photons to characterize the steady-state fluorescent background and fit a bi-exponential decay model to retrieve two fluorescence lifetime components, 1.40±0.27 and 0.46±0.02 ns, with amplitudes of 21.80±2.20, and 0.30±0.20, respectively (Fig. 2e, Supplementary Fig. 2a and Table 1). The mean fluorescence lifetime of the background photons (1.39±0.27 ns) closely mirrors the primary long lifetime component of the bi-exponential decay. Additionally, the fraction of the long lifetime component is 0.986±0.009.

**Table 1:** Best-fit results of tail-fitting fluorescence decays with sum of exponential decay models.

| Photon class | $\tau_1$ (ns)<br>(for $n=2$)<sup>1</sup> | $\tau_2$ (ns)<br>(for $n=2$) | $a_1$<br>(for $n=2$) | $a_2$<br>(for $n=2$) | $\tau$ (ns)<br>(for $n=1$)<sup>2</sup><br>/<br>$\bar{\tau}$ (ns)<br>(for $n=2$)<sup>3</sup> | $f_1$ (for $n=2$)<sup>4</sup> |
| --- | --- | --- | --- | --- | --- | --- |
| Diffused mCherry inside precipitates | 1.54±0.01<sup>5</sup> | 0.55±0.01<sup>6</sup> | 14.4±0.2<sup>7</sup> | 86.6±1.4<sup>8</sup> | 0.86±0.01<sup>9</sup> | 0.142±0.007<sup>10</sup> |
| mCherry clusters inside precipitates |  |  |  |  | 1.07±0.35 |  |
| Diffuse mCherry-HP1α inside biocondensates | 1.40±0.27 | 0.46±0.02 | 21.80±2.20 | 0.30±0.20 | 1.39±0.27 | 0.986±0.009 |
| Diffuse mCherry-HP1α outside biocondensates | 1.41±0.03 | 0.46±0.00 | 32.12±0.85 | 0.70±0.01 | 1.41±0.03 | 0.998±0.001 |
| mCherry-HP1α clusters in biocondensates in ESCs | - | - | - | - | 1.20±0.15 | - |
| mCherry-HP1α clusters in biocondensates in 2d-RA ESCs | - | - | - | - | 0.82±0.17 | - |
| Diffuse mCherry-CENP-V inside biocondensates | 1.19±0.01 | 0.18±0.00 | 28.80±3.32 | 1.15±0.30 | 1.18±0.01 | 0.961±0.011 |
| Diffuse mCherry-CENP-V outside biocondensates | 1.42±0.01 | 0.25±0.00 | 50.85±1.35 | 0.25±0.14 | 1.42±0.01 | 0.995±0.002 |
| mCherry-CENP-V clusters in biocondensates in ESCs | - | - | - | - | 1.18±0.22 | - |
| Diffuse mCherry-NCL inside biocondensates | 1.32±0.01 | 0.19±0.00 | 20.22±2.36 | 0.57±0.03 | 1.31±0.01 | 0.972±0.008 |
| Diffuse mCherry-NCL outside biocondensates | 1.39±0.01 | 0.25±0.00 | 30.65±0.28 | 0.12±0.00 | 1.39±0.01 | 0.996±0.001 |
| mCherry-NCL clusters in biocondensates in ESCs | - | - | - | - | 1.20±0.38 | - |

<sup>1</sup>Two exponential decay components in using eq. 1 (see Methods) for tail-fitting fluorescence decays
<sup>2</sup>One exponential decay component in using eq. 1 (see Methods) for tail-fitting fluorescence decays
<sup>3</sup>Calculated using eq. 2 (see Methods) for a bi-exponential fluorescence decay model
<sup>4</sup>Calculated using eq. 3 (see Methods) for a bi-exponential fluorescence decay model
<sup>5,6,7,8</sup>Fitting standard errors of a bi-exponential decay model parameters
<sup>9</sup>If derived from mono-exponential decay model, then error value is the fitting standard errors of a mono-exponential decay model parameters. If derived as mean fluorescence lifetime from a bi-exponential decay model, then error value was propagated from employing eq. 4 on the function in eq. 2 (see Methods)
<sup>10</sup>Error value was propagated from employing eq. 4 on the function in eq. 3 (see Methods)

This suggests a majority fraction of diffuse mCherry-HP1α molecules inside the biocondensate, which are not sufficiently bright to show up as photon bursts and have fluorescence lifetimes slightly smaller than the natural fluorescence lifetime of mCherry in densities <30% fractional volume occupancy (FVO)^45^. Next, as a control, we examine the fluorescence lifetimes of mCherry-HP1α molecules outside biocondensates in both 2d-RA and undifferentiated ESCs together. Again, we fit a bi-exponential decay model and retrieve two fluorescence lifetime components, 1.41±0.03 and 0.46±0.01 ns, with amplitudes of 32.12±0.85, and 0.70±0.01, respectively (Supplementary Fig. 2b and Table 1). This yields a mean fluorescence lifetime of 1.40±0.03 ns, which again closely mirrors the primary long lifetime component. Additionally, the fraction of the long lifetime component is 0.998±0.001. This suggests that mCherry-HP1α outside of biocondensates also have fluorescence lifetimes slightly smaller than the natural fluorescence lifetime of mCherry in densities <30% FVO^45^. Next, we employ the burst search and selection algorithm. In this filtering process, we exclude the diffuse fraction as well as bursts with insufficient photon rate for fluorescence lifetime analysis and focus on fluorescence originating from large clusters of mCherry-HP1α, which yield a fluorescent burst with a photon rate that is at least six times higher than the background photon rate. From a total of 45 different cells and ROIs per condition, we filtered a total of 60 and 11 bursts across all undifferentiated and 2d-RA ESCs samples, respectively (Fig. 2f). This leads to an overall burst detection rate of 1.33 and 0.24 bursts per ROI in undifferentiated and 2d-RA ESCs, respectively. While we find a sufficient number of these clusters in undifferentiated ESCs, the few clusters found in 2d-RA ESCs are not statistically sufficient to consider their size, brightness or lifetime distributions.

Secondly, we fit the combined fluorescence decay of all burst photons from each condition to a mono-exponential decay function. We find fluorescence lifetimes of 1.20±0.15 and 0.82±0.17 ns for mCherry-HP1α clusters found in undifferentiated and 2d-RA ESCs, respectively (Supplementary Fig. 2c, d, and Table 1). Importantly, while both decays are noisy, since we do not acquire as many bright bursts from samples at 2d-RA condition, its model-fitting results are unreliable (Supplementary Fig. 2d). Overall, in undifferentiated ESCs, there are more clusters of mCherry-HP1α, with mean fluorescence lifetimes substantially lower (1.20±0.15 ns) than in diffuse mCherry-HP1α (1.39±0.27 ns). In undifferentiated ESCs, mCherry-HP1α clusters are denser, showing a further reduction in lifetime compared to the steady-state background.

Significantly, mCherry-HP1α clusters in chromocenters of undifferentiated ESCs show narrow fluorescence lifetime distribution with a mean lower by ∼0.3 ns than the mean lifetime value we recover using FLIM (Fig. 2i-k). The burst width analysis (Fig. 2g), as well as fluorescence correlation spectroscopy analysis (FCS)^59^ (Supplementary Fig. 3a-b) suggest these mCherry-HP1α clusters exhibit few ms diffusion through the confocal volume (Supplementary Table 2). These fractions, mainly clusters, recur in heterochromatin condensates in ESCs and, to a much lesser extent, in 2d-RA ESCs. Recent studies show direct link between the fluorescence lifetime reduction of mCherry and the local density in its vicinity^45,60^. Based on these findings, our results suggest that mCherry-HP1α clusters are denser than diffuse mCherry-HP1α in chromocenters of ESCs.

As a control to test whether BLISS indeed detects clusters of mCherry-HP1α recurring into the photobleached ROI from outside of it, we challenged the experiment against the same conditions only after cell fixation using paraformaldehyde (Fig. 3a, see Methods), hence in chromocenters that are expected to be largely immobile. Firstly, we retrieve steady-state background rates starting from 2.0 and ending at 0.8 kHz, for mCherry-HP1α measured in fixed undifferentiated ESCs during continuous photobleaching (Fig. 3b, Supplementary Fig. 1c). We acquire the fluorescence decays accumulated from photons detected outside bursts as part of the steady-state fluorescent background and fit a bi-exponential decay model to retrieve two fluorescence lifetime components, 1.66±0.01 and 0.47±0.01 ns, with amplitudes of 1.47±0.02 and 1.59±0.05, respectively (Supplementary Fig. 4a). The mean fluorescence lifetime of the steady-state background photons is 1.38±0.01 ns, similar to the lifetime of the steady-state background in unfixed undifferentiated ESCs. Next, as a control, we examine the fluorescence lifetimes of mCherry-HP1α molecules outside biocondensates. Again, we fit a bi-exponential decay model and retrieve two fluorescence lifetime components, 1.61±0.01 and 0.32±0.00 ns, with amplitudes of 8.9±0.1, and 5.0±0.1, respectively (Supplementary Fig. 4b), yielding a mean fluorescence lifetime of 1.48±0.01 ns, which is the natural fluorescence lifetime of mCherry. Unlike with living ESCs, here we only find eight bursts inside chromocenters from a total of 22 BLISS measurements. This leads to an overall burst detection rate of 0.36 bursts per ROI, hence the rate of detection was reduced by 97% compared to the living ESCs samples. Measuring FCS inside chromocenters of fixed undifferentiated ESCs, shows that we brought diffusion to a halt (Supplementary Fig. 3c), suggesting that the very few bursts found in fixed ESCs are potentially due to rare deviations of a noisy signal from the mean of the fluorescence background, and not due to actual diffusing clusters.

**Figure 3.**
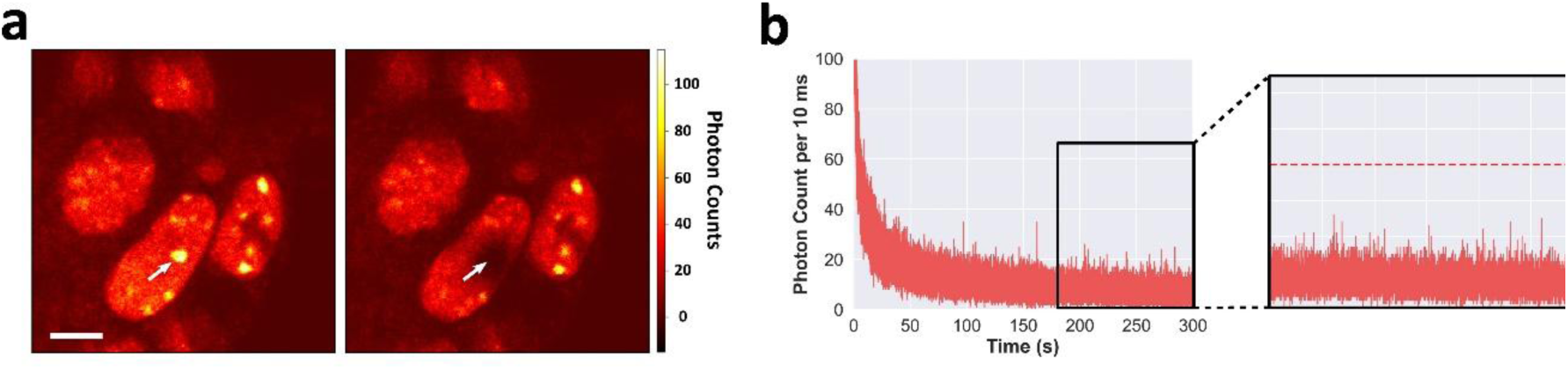
BLISS performed in fixed undifferentiated mouse ESCs expressing mCherry-HP1α shows significant decrease in recurring HP1α clusters. (a) Images of cell nuclei, focusing on a ROI (white arrows) in one nucleus before (left) and after (right) continuous photobleaching and collection of fluorescence from fixed undifferentiated mouse ESCs. (b) Example of 300 s fluorescence time trace of fixed undifferentiated mouse ESCs, that exhibit continuous photobleaching until reaching a steady-state with low counts serving as background levels, atop which photon bursts do not appear. A minimal SBR threshold of F=6 (indicated as red dashed line), is used for identifying photon bursts and minimizing false detection of noise fluctuations about the mean fluorescence. Scalebar 5 μm.

### BLISS recovers CENP-V and nucleolin dynamic clusters with densities similar to those recovered from FLIM

HP1α is not the only constituent of heterochromatin condensates and of chromocenters particularly. To extend BLISS measurements in chromocenters beyond HP1α, we selected an additional chromocenter-residing protein, namely the kinetochore protein Centromeric protein V (CENP-V), from our endogenously-tagged mCherry ESC library^44^. CENP-V is known to interact with HP1α and is an important linker of the primary constriction of chromocenters^61^. We measured single clusters of mCherry-CENP-V recurring inside chromocenters of undifferentiated ESCs during its continuous photobleaching. Then, using the same approach, we analyze the fluorescence lifetime of photon bursts arising from these clusters. First, we retrieve a steady-state background rate of approximately 1.0 kHz (Supplementary Fig. 1c). As control, we examine the fluorescence lifetimes of mCherry-CENP-V molecules outside chromocenters. We fit a bi-exponential decay model and retrieve two fluorescence lifetime components, 1.42±0.01 and 0.25±0.00 ns (Supplementary Fig. 5a), with a mean fluorescence lifetime of 1.42±0.01 ns since the fraction of the longer fluorescence lifetime is 0.995±0.002. Next, we employ the burst search and selection algorithm to select a total of 41 bursts from 37 different cells and ROIs giving a detection rate of 0.90 bursts per ROI. Fitting the combined fluorescence decay of all burst photons to a mono-exponential decay model yields a mean fluorescence lifetime of 1.18±0.22 ns (Supplementary Fig. 5b and Table 1). These results suggest that mCherry-CENP-V clusters inside chromocenters have a mean fluorescence lifetime that is lower (1.18±0.22) than diffuse mCherry-CENP-V (1.41±0.01). Pixel-wise fluorescence lifetime distribution recovered from FLIM of ESCs expressing mCherry-CENP-V, shows a mean fluorescence lifetime of 1.24 ns (Fig. 4a-b, e). Similarly, the fluorescence lifetime distribution of 41 bursts retrieved using BLISS gives a mean value of 1.27 ns (Fig. 4f). These results suggest that CENP-V clusters in chromocenters of undifferentiated ESCs have similar density to their pixel-wise average density.

**Figure 4.**
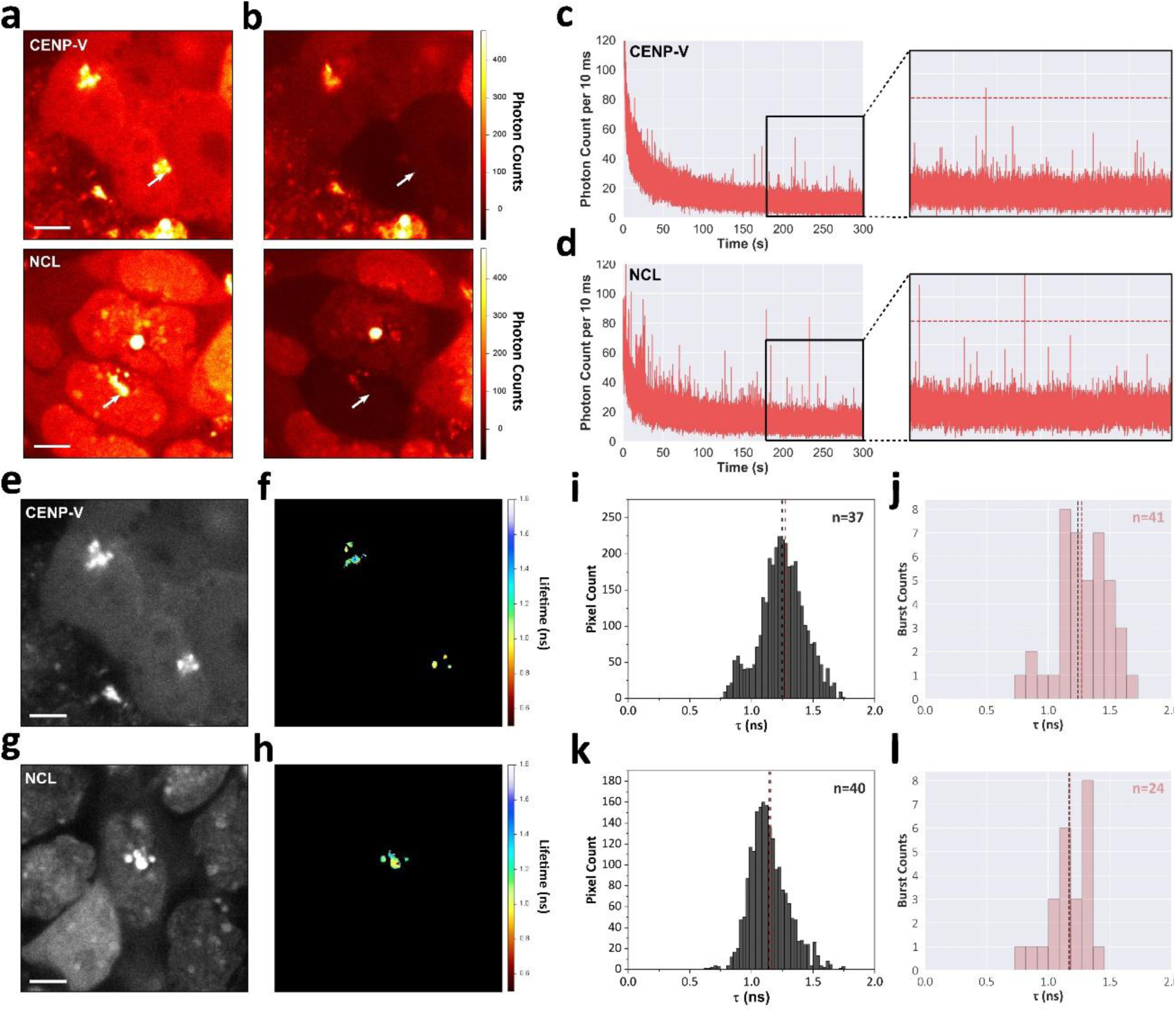
BLISS on mouse ESCs expressing mCherry-CENP-V and mCherry-NCL recovers clusters of CENP-V or NCL of similar densities to the mean density recovered from FLIM. (a-b) Fluorescence intensity images of cell nuclei, focusing on a ROI (white arrows) in one nucleus before (a) and after (b) continuous photobleaching and collection of fluorescence, of mouse ESCs expressing mCherry-CENP-V (top) or mCherry-NCL (bottom). (c-d) Examples of 300 s fluorescence time traces of a ROI in mouse ESCs nuclei expressing mCherry-CENP-V (c) or mCherry-NCL (d). The time traces exhibit continuous photobleaching, until reaching a steady-state with low counts serving as background levels, atop which photon bursts clearly appear. A minimal single-to-background ratio (SBR) threshold of F=6 (indicated as red dashed line), is used for identifying these photon bursts and minimizing false detection of noise fluctuations about the mean fluorescence. (e, g) Example intensity images and FLIM (f, h) of mouse ESCs expressing mCherry-CENP-V (e, f) or mCherry-NCL (g, h). (i, k) Pixel-wise mean fluorescence lifetime histograms recovered from FLIM of ESCs expressing mCherry-CENP-V (i) or mCherry-NCL (k). (j, l) Burst-wise mean fluorescence lifetime histograms recovered from BLISS of mouse ESCs expressing mCherry-CENP-V (j) or mCherry-NCL (l). Black dashed line – mean fluorescence lifetime recovered from FLIM. Red dashed line – mean fluorescence lifetime recovered from BLISS. Scalebars, 5 μm.

We next sought to apply BLISS to a non-chromocenter protein, but which nonetheless participates in nuclear biocondensates. We once again turned to our endogenously-tagged mCherry library^44^, and selected the nucleolar protein nucleolin (NCL). As in the previous examples, we employ FLIM to recover pixel-wise mean fluorescence lifetimes and BLISS to recover fluorescence lifetime distribution arising from clusters. First, we retrieve a steady-state background rate of approximately 0.5 kHz (Supplementary Fig. 1d). As control, the fluorescence lifetimes of mCherry-NCL molecules outside of their relevant biocondensates are 1.39±0.01 and 0.25±0.00 (Supplementary Fig. 5c). With the fraction of the longer lifetime being 0.996±0.001, the calculated mean fluorescence lifetime is 1.39±0.01 ns. Next, we employ the burst search and selection algorithm to select a total of 24 bursts from 39 different cells and ROIs, giving a detection rate of 0.61 bursts per ROI. Fitting the combined fluorescence decay to a mono-exponential decay model yields a mean fluorescence lifetime of 1.20±0.38 ns (Supplementary Fig. 5d and Table 1). This suggests that mCherry-NCL clusters inside nucleoli are denser than diffuse mCherry-NCL. Pixel-wise fluorescence lifetime distribution recovered from FLIM of undifferentiated ESCs expressing mCherry-NCL yield a mean value of 1.14 ns (Fig. 4c-d, g). Using BLISS, we find 24 bursts arising from mCherry-NCL clusters and recover a single population distribution of fluorescence lifetimes with a mean value of 1.15 ns (Fig. 4h). Based on these findings, much like mCherry-CENP-V, also mCherry-NCL forms clusters that recur inside its relevant biocondensates, with fluorescence lifetime distributions that are similar to the fluorescence lifetimes recovered by FLIM. This suggests that NCL clusters have similar density to their pixel-wise average density. Next, we employ FCS analysis on mCherry-CENP-V and mCherry-NCL in ESCs (Supplementary Fig. 3d, e). We find that while mCherry-CENP-V exhibits tens-of-ms diffusion time through the confocal volume, mCherry-NCL exhibits hundreds-of-ms diffusion time in similar conditions (Supplementary Table 2).

## Discussion

The study of biomolecular condensates and heterochromatin domains using standard confocal microscopy relies on pixel-level ensemble-averaged imaging, which inherently masks local sub-populations and transient micro-architectures. By introducing BLISS, we demonstrate that a standard laser-scanning confocal setup can bypass pixel-level ensemble averaging to resolve individual, rare dynamic biomolecular clusters within dense cellular ROIs. Our approach reveals that some nuclear condensates are not uniform phase-separated droplets, but rather display a compartmentalized network harboring distinct, hyperdense heterogenous nanoscale assemblies that dynamically exchange depending on their immediate environment.

Applying BLISS to chromocenter in mouse ESCs uncovers a previously hidden sub-population of mCherry-HP1α clusters possessing significantly lower fluorescence lifetimes (0.9-1.1 ns) than the pixel-level ensemble average (∼1.3 ns). Because mCherry fluorescence lifetime decreases predictably with increasing local density via a self-quenching mechanism^45^, these shorter lifetimes directly map to local hyperdense micro-domains^57,58^ (Fig. 5). Crucially, on the one hand, these hyperdense HP1α assemblies are hyperdynamic, since they diffuse into the photobleached ROI on millisecond timescale through a tightly focused confocal volume and are predominantly restricted to undifferentiated mouse ESCs^38,62–65^. Yet, in 2d-RA ESCs the number and rate of these dynamic HP1α assemblies reduce substantially as chromocenters become more homogenous^45^ (Fig. 5). This dramatic shift points to fundamental biophysical mechanism underlying pluripotency; in stem cells, heterochromatin is maintained in a globally decondensed high plasticity state by a mobile pool of loosely-bound hyperdynamic HP1α assemblies^38,62–65^ (Fig. 5). As cells commit to a lineage, this hyperdynamic fraction consolidates, driving heterochromatin maturation into a more rigid, homogenous architecture^45^.

**Figure 5.**
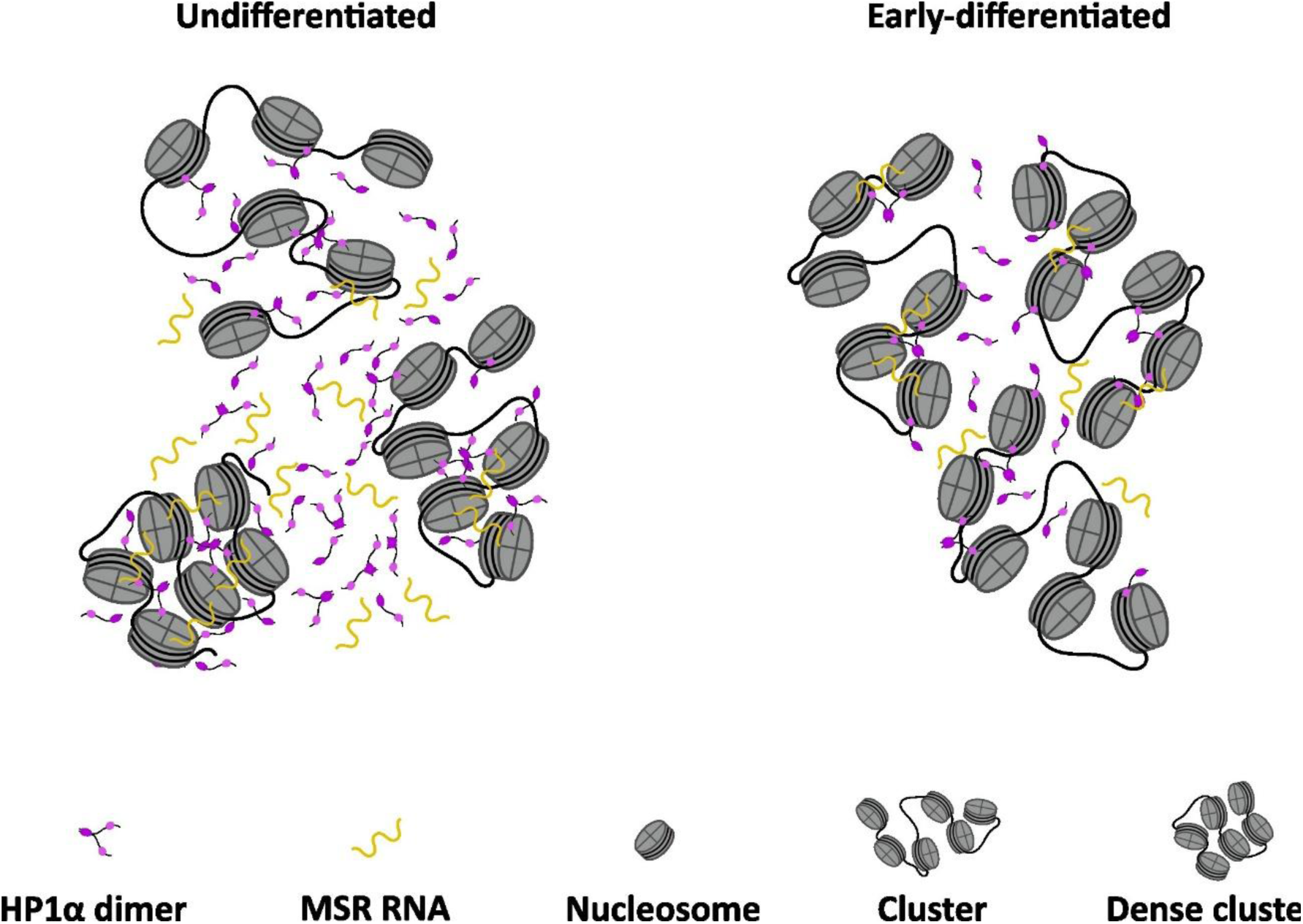
Illustration of HP1α nucleosome clusters and their differences between undifferentiated and early-differentiated mouse ESCs. In undifferentiated cells clusters are found in different densities and are sparse due to low abundance of H1. In early-differentiated cells clusters are less dense and more homogeneous due to higher concentration of the H1 linker histone.

BLISS thus provides a direct quantitative readout of chromatin liquidity during cellular differentiation. Importantly, our findings demonstrate that local hyper density is not a universal artifact of all condensates-associated proteins. While kinetochore protein CENP-V and the nucleolar protein NCL also form recurring dynamic clusters under BLISS, their single-cluster fluorescence lifetime distribution closely matches their pixel-level ensemble-averaged FLIM values (∼1.30 and ∼1.15 ns, respectively). This indicates that the local density of CENP-V and NCL within BLISS detectable bursts is similar to the mean density of their phase-separated condensates. This stark contrast between HP1α and other nuclear scaffolders suggests structural specificity; whereas CENP-V and NCL partition homogeneously throughout their respective domains, while HP1α forms multi-valent, higher-order topological networks likely bridged by MSR RNA^45,47–49^, linker histone H1^46^ and other potential interactors bridging nucleosome clusters and generating localized high-density hubs. The lower abundance of linker histone H1 in pluripotent ESCs relative to early-differentiated ESCs^66^ is consistent with the heterogeneous chromatin density observed in pluripotency, whereas higher H1 levels correlate with structural homogenization.

Methodologically, BLISS bridges a critical gap between test-tube single-molecule spectroscopy and live-cell super-resolution imaging. Conventional single-molecule methods in dense regions typically require ultra-low expression levels^3,21,23,24^, photo-switchable fluorophores^5–7^, specialized illumination geometries (TIRF/HILO)^33–37^, or expensive super-resolution optics^10,11,14,16,17^. By harnessing continuous photobleaching to establish a controlled steady-state background, BLISS achieves single-particle sensitivity using endogenous FP tags and conventional confocal microscopy setups already available in most life-science institutes. Importantly, the fluorescent tags under investigation need to withstand continuous photobleaching sufficiently to yield a burst of photons with photon rate significantly above the steady-state background rate.

While BLISS relies on sufficient brightness contrast and steady-state diffusion balance, its operational principles extend far beyond heterochromatin. This framework can be readily adapted to investigate transcription factor clustering at super-enhancers^67^, RNA Pol II hubs^68^, synaptic density assemblies^69^, and DNA damage repair foci^70^. Future integration of multi-color BLISS will allow researchers to perform burst-wise Förster resonance energy transfer (FRET) efficiency^1,71^ and alternating laser excitation (ALEX) / pulsed interleaved excitation (PIE) stoichiometry^72,73^, offering a generalizable tool.

## Methods

### PyBroMo simulations

Simulations of single-molecule photon bursts in the presence of a high concentration of background diffuse molecules were achieved using PyBroMo python-based simulations approach^74,75^. These simulations tested different conditions mimicking reaching a steady state after completing the fluorescence reduction in continuous photobleaching. PyBroMo simulations are performed in two steps: (i) simulating the diffusion trajectory of each simulated molecule within a box of given dimensions, knowing the diffusion coefficient of each molecule, and then (ii) based on a PSF function in space, the emission rate of each simulated molecule was calculated depending on where it was relative to the center of the PSF at each moment of the simulation. This resulted in photon bursts arising from sufficiently bright molecules when crossing the central part of the PSF, which is equivalent to the confocal volume. After assigning spatial positions for each of the simulated molecules, in each 50 μs iteration time, with simulation times lasting ten minutes, we calculated the photon rate of each molecule in each position along the simulation and summed the rates for each instance. Then, FRETbursts was used to (i) estimate the background rate, and (ii) search and select photon bursts (see data analysis in *Microscopy Experiments and Analyses* section), based on different minimal SBR thresholds. The settings of the simulations and the results of burst analyses are provided in Supplementary Table 1. In these simulations, two subpopulations of diffusing species were simulated: (i) a large number of rapidly-diffusing molecules with low molecular brightness, yielding the fluorescent background reached at steady-state following continuous photobleaching, and (ii) a small number of slowly-diffusing molecules with high molecular brightness, yielding the photon bursts atop the fluorescent background. We controlled the concentration of the two subpopulations of molecules by dividing the number of molecules from that subpopulation by the volume of the simulation box. Of note, we strive to reach a few tens of pM for the bright molecules, to match typical confocal-based single-molecule spectroscopy concentrations. The detector’s typical background was taken as 200 Hz, which is relevant to the hybrid photomultiplier tubes (PMTs) we use in our experimental setup.

### Microscopy Experiments Setup and Analyses

FLIM and BLISS measurements of mCherry-HP1α in mouse ESCs were performed using a laser scanning confocal microscopy setup with time-correlated single photon counting (TCSPC) time-resolved capabilities, as was described previously^45^. In BLISS experiments, the confocal volume was brought to a selected pixel within an acquired frame and parked there for a 5-minute data acquisition, for each technical repeat. The power of the 532 nm pulsed laser (20 MHz repetition rate, and pulse FWHM of 150 ps) that was used in BLISS experiments, to achieve both efficient photobleaching and high photon rates from bursts, was 125 μW. Importantly, a FLIM image was taken before and after the BLISS data acquisition.

The files of each 5-minute data acquisition were converted to the photon-HDF5 file format^56^, and were then analyzed to search and select photon bursts, using the FRETbursts software^50^. Firstly, the background rate was estimated for every 40 s of the measurements. Then, knowing the changing background rate, bursts were searched using a sliding window of m (=10) consecutive photons, seeking instances in which the instantaneous photon rate for these m-photons was at least F (=6) times higher than the background rate, setting a minimal SBR of F-1 (=5)^76^. A burst consists of a collection of consecutive photon detections, within which every subset of m consecutive photons had an instantaneous photon rate higher than F times the background rate. This results in bursts with sizes (i.e., total number of photons) of at least 10 photons, which would be insufficient for accurate calculation of the burst fluorescence lifetime from the mean of all photon nanotimes, where a nanotime is the time, a photon was detected relative to the laser pulse moment that led to its detection. Therefore, bursts were selected with a minimal threshold on the burst sizes of 40 photons. Then, different burst-wise parameters were calculated and presented in histograms for all acquired bursts in all measurements of the same conditions. These parameters include (i) burst sizes, (ii) burst widths, which are the duration of diffusion through the confocal volume, (iii) maximal instantaneous photon rate, which is equivalent to the molecular brightness, and (iv) mean photon nanotimes.

Using the same burst search algorithm, we extracted photons outside of bursts by subtracting all bursts found using F=2 from those found using F=1.

Fluorescence decays of different classes of detected photons were formed from the histogram of photon detection nanotimes. Then, fluorescence decays have undergone tail-fitting to sum of *n* exponential decays (eq. 1), whether to mono-exponential decays (*n*=1) or to bi-exponential decay (*n*=2) models, where the model choice was the one that included the minimal number of free parameters that are statistically sufficient to describe the experimental fluorescence decay. For that matter, the fluorescence decay of photons outside bursts was clearly not mono-exponential and hence has undergone tail-fitting to a bi-exponential decay model. Fluorescence decays of burst photons included much less photons, and a mono-exponential decay model was sufficient for tail-fitting.

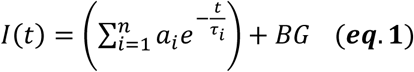

Whenever fluorescence decays models of more than a single exponential decay component were used, the mean fluorescence lifetime (eq. 2) and the fraction of the *i*^th^ fluorescence lifetime component (eq. 3) were calculated from the best-fit parameter values from fitting a bi-exponential decay function.

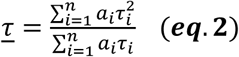

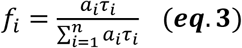

The errors of values, *δP*, of parameters, *P*, calculated from eqs. 2, 3, were propagated from the standard errors of best fit parameter values, *δp_i_* (eq. 4).

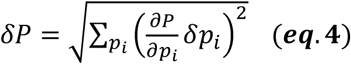

FCS data was acquired from the steady-state fluorescence of BLISS measurements. The autocorrelation function was calculated from the fluorescence trajectories. The fluorescence autocorrelation functions were then fitted to a model with a translational 3D diffusion process and an exponential rapid dynamic process (eq. 5):

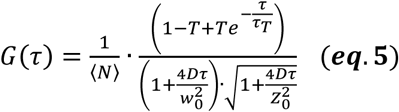

where *τ* is the lag time, *G*(*τ*) is the autocorrelation of the data at different lag times, ⟨*N*⟩ is the mean number of fluorescent molecules in the effective excitation volume, *D* is the translational diffusion coefficient, *w*_0_ and *z*_0_ are the width and height of the effective excitation volume, respectively. *τ_T_* and *T* are the relaxation time and the equilibrium fraction, respectively, of a dim state of a fluorescence fluctuation process between bright and dim states (see eq. 6).

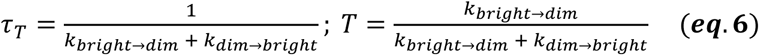

To calculate the diffusion time (*τ_D_*), we use the following (eq. 7):

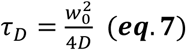

### Sample Preparation

Mouse ESCs were prepared as described in Joron, Viegas *et al.*^45^, under the following conditions. Mouse ESCs were cultured using standard procedures on mouse embryonic fibroblast (MEF) on 0.2% gelatine coated plates, in ESC medium based on Dulbecco’s modified eagle medium (DMEM; Sigma) supplemented with 15% fetal bovine serum (FBS; Biological Industries), L-glutamine, 1X minimum essential medium (MEM) non-essential amino acids, Sodium Pyruvate, 1x Penstrep, 0.2 mM β-ME and Leukemia inhibitory factor (LIF). Cell cultures were maintained in a humidified atmosphere (5% CO_2_ at 37 °C). Depletion of MEFs was performed by pre-plating trypsinized ESCs on low attachment plates twice, for 30 mins each. For measurements at *2i* conditions, ESCs were cultured without MEFs on 0.2% gelatin coated plates for a few passages in the ESC medium with only 2% FBS and with the MEK inhibitor, Mirdametinib (PD0325901) and the GSK-3 inhibitor and Wnt/β-catenin activator CT-99021 (CHIR99021), together with LIF. For early differentiation measurements, cells were grown for 3 days on 0.2% gelatin coated plates without MEFs with ESC medium without LIF and with 1 µM RA. For live imaging, cells were seeded on an ibiTreat 8-well µ-slide (#80826 Ibidi, Munich, Germany) in appropriate cell culture medium. For fixed cell imaging, cells were fixed in 4% paraformaldehyde (PFA) at room temperature (25°C) for two rounds. First, the cells were incubated for 15 minutes, then washed with PBS buffer 3 times and left overnight in PBS at 4°C. In the second round, the cells were incubated for 42 minutes in 4% PFA and washed again 3 times with PBS, then mounted using Fluoromount-G^®^ (SounthernBiotech).

## Acknowledgements

This work was supported by the Israel Science Foundation (ISF) (556/22 to E.L.), ISF’s Personalized Medicine Award (IPMP) (3605/21 to E.M.), The Israel Ministry of Science (MOST 0004272 to E.M.), the European Union’s Horizon Europe Research and Innovation Programme under the EIC Pathfinder-Open grant agreement #101099654 (RT-SuperES), and by Volkswagen Foundation (0200195-03 to E.L.). E.M. is the incumbent of the Arthur Gutterman Family Chair for Stem Cell Research. K.J. is supported by the Clore Israel Foundation.

## Author Contributions

K.J. and E.L. conceived the methodological concept; K.J. and E.M.1 designed and performed all experiments; K.J. analyzed the data; E.L. supervised the study and helped with data analysis; E.M.2 designed the experiments and supervised the study. K.J., E.M.1, E.M.2, and E.L. wrote the paper.

## Data Availability

All simulated and experimentally-acquired data have been deposited on Zenodo [https://doi.org/10.5281/zenodo.22079288]. Additionally, the whole analytical pipeline has been summarized in a Jupyter notebook, deposited over Zenodo [https://doi.org/10.5281/zenodo.22079288].

