## Supplementary Information for "BLeaching In-cell Single-molecule burstS (BLISS) reveals dynamic fraction of HP1α clusters in chromocenters of undifferentiated embryonic stem cells"

Khalil Joron[1], Eden Mishne[2], Eran Meshorer[2,3], Eitan Lerner[1,4]

[1] Department of Biological Chemistry, The Alexander Silberman Institute of Life Sciences, Faculty of Sciences, The Edmond J. Safra Campus, The Hebrew University of Jerusalem, 9190401, Jerusalem, Israel.

[2] Department of Genetics, The Alexander Silberman Institute of Life Sciences, Faculty of Sciences, The Edmond J. Safra Campus, The Hebrew University of Jerusalem, Jerusalem, 9190401, Israel.

[3] Edmond and Lily Center for Brain Sciences (ELSC), The Edmond J. Safra Campus, The Hebrew University of Jerusalem, Jerusalem, 9190401, Israel

[4] The Center for Nanoscience and Nanotechnology, The Edmond J. Safra Campus, The Hebrew University of Jerusalem, Jerusalem, 9190401, Israel.

### Supplementary information includes:

- Supplementary text: detailed explanation of the PyBroMo simulations
- Supplementary figures 1-5
- Supplementary tables 1-2

### Supplementary text

*Simulations of rare clusters in presence of predominant diffuse molecules in BLISS*

In the presented experimental approach, we use a pulsed laser focused into the cell at a given ROI, operating at high power (~125 μW) for a given amount of time, and fluorescence photons are detected during the course of continuous photobleaching. The typical non-fluorescent background rates of the detectors are known and can be compared to the counts-per-second levels at each moment along the course of the continuous photobleaching measurement. After an initial time window, during which the counts-per-seconds levels dropped and stabilized close to steady-state, we search for photon bursts using the common sliding window burst search approach. In short, this is achieved by using a sliding window of *m* consecutive photons to estimate the instantaneous photon rate. A burst is the collection of consecutive photons, where each *m*-photon time window has an instantaneous rate ≥*F* times the temporal background rate. Then all identified photon bursts have burst parameters, such as width, defined by the interval between the first and last photon detection time, and burst size, defined by the total number of photons. Importantly, the higher the minimal instantaneous rate threshold, *F*, is, the lower the probability that a fluctuation of the background above the mean would falsely be detected as a signal. Typically, using *F=6* (minimal SBR of 5) sufficiently suppresses the detection of background fluctuations. Moreover, the higher the value of *F* is, the closer to the center of the confocal volume is the underlying diffusing fluorescent particle from which burst photons originate[55]. After identifying photon bursts, they are further filtered based on thresholds of burst parameter values, such as a burst size threshold. For the purpose of calculating burst-wise mean photon detection nanotimes (times relative to moment of pulsed excitation) with values that accurately map to mean fluorescence lifetimes, a minimum of ~40-50 photons/burst detection channel is required. Considering the herein presented approach, data attained in BLISS experiments can be simulated using simulations of freely-diffusing fluorescent particles near a confocal volume.

We use the PyBroMo python-based simulations approach[76,77]. We simulate diffusion of many rapidly diffusing molecules with low brightness together with a few slowly-diffusing brighter molecules, testing the capability to quantify background rates and photo bursts in different scenarios. We test suboptimal conditions using a 10-fold ratio between the brightnesses of the two diffusing species.

Then, bursts were identified from the simulation results, with the aim to (i) maximize the number of bursts with sizes ≥50 photons for accurate mean fluorescence lifetime calculation per burst, and to (ii) maximize the number of bursts arising from the slow diffusing clusters. The results of the simulations are shown in Supplementary table 1. In simulations in which the steady-state concentration of rapidly diffusing low brightness molecules was 3.1 nM, instantaneous photon rate thresholds, *F*, of 6 or higher yielded hundreds of bursts, with a large portion arising from the slow diffusing clusters. Of note, the larger the brightness ratio was between the two diffusing molecular species, the more bursts were identified with more of them arising from the slow diffusing clusters. If, however, continuous photobleaching led to a lower steady-state concentration of rapidly diffusing low brightness molecules, such as 0.5 nM, then less bursts arise from the slow diffusing clusters, and more of the rapidly diffusing molecules show up, not only as a background but also as bursts.

Based on these results a few guidelines can be drawn for resolving bursts of slow diffusing bright species in the presence of many rapidly diffusing low brightness species. First, the steady-state concentration of fluorescent molecules depends on the laser intensity and on the initial concentration before the experiment started. Therefore, the identification of bursts arising mostly from the rare subpopulation of slow diffusing clusters strongly depends on the laser intensity used. Second, the higher the ratio of the brightness of the diffusing species is, the easier it would be to detect bursts arising mostly from the slow diffusing bright species. Third, in burst search, instantaneous photon rate thresholds are recommended to be ≥6.

### Supplementary figures

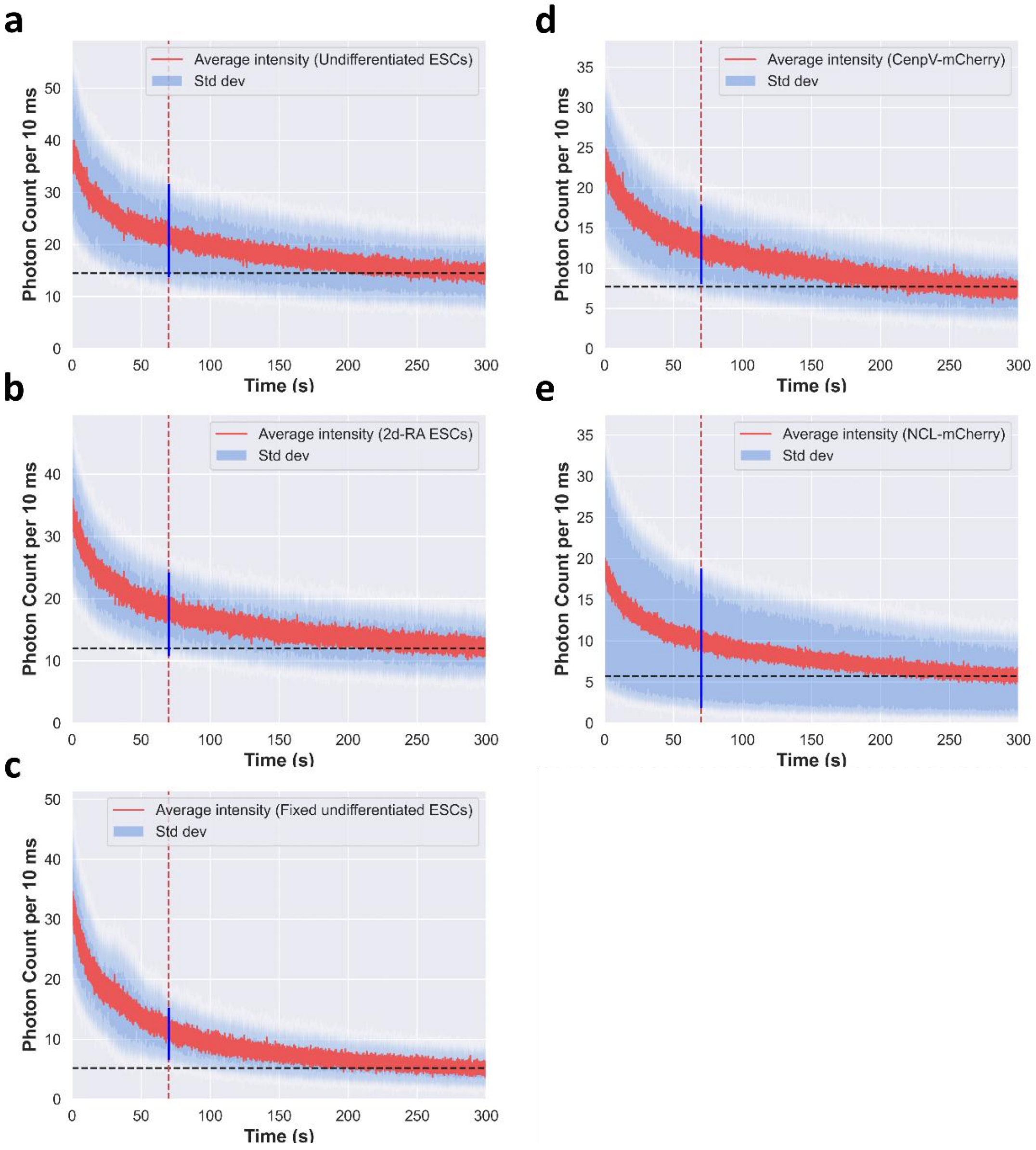


**Supplementary figure 1. Mean fluorescence time traces of all BLISS technical repeats of each condition.** Combined fluorescence time-trace of mCherry-HP1α in pluripotent undifferentiated (a), early-differentiated (b) and fixed undifferentiated mouse ESCs (c). Combined fluorescence time-trace of mCherry-CENPV (d) and mCherry-nucleolin (e). The max width of values of photon count rates is 8, 12 or 16 counts/10 ms, for fixed-undifferentiated, 2d-RA and undifferentiated ESCs, respectively, and 16 or 3 for CENP-V and nucleolin, respectively, indicated by the highlighted blue line atop the curves. Black dash line indicates the average steady-state photon count reached at the end of measurement, which is 15, 12 and 5 counts/10 ms for undifferentiated, 2d-RA and fixed-undifferentiated mouse ESCs, respectively, and 8 or 6 for CENP-V and nucleolin, respectively.

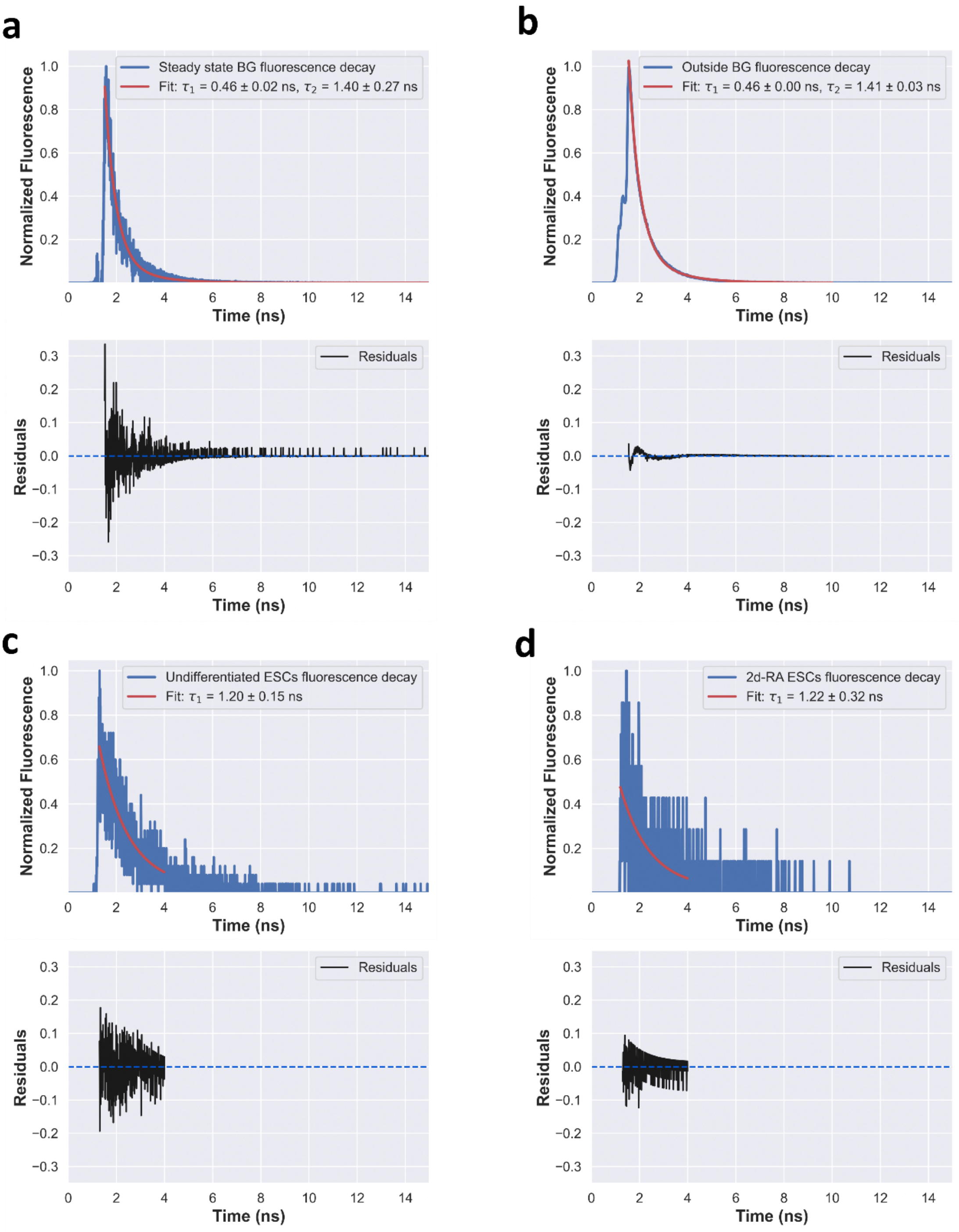


**Supplementary figure 2. Fitting results to fluorescence decays of mCherry-HP1α.** The fluorescence decay curves and the fitting results, including residuals, of mCherry-HP1α from inside (a) and outside (b) heterochromatin biocondensates after reaching steady-state background. The fluorescence decay curves and fitting results, including residuals, of recovered mCherry-HP1α fluorescent bursts inside heterochromatin biocondensates of undifferentiated ESCs (c) and 2d-RA ESCs (d).

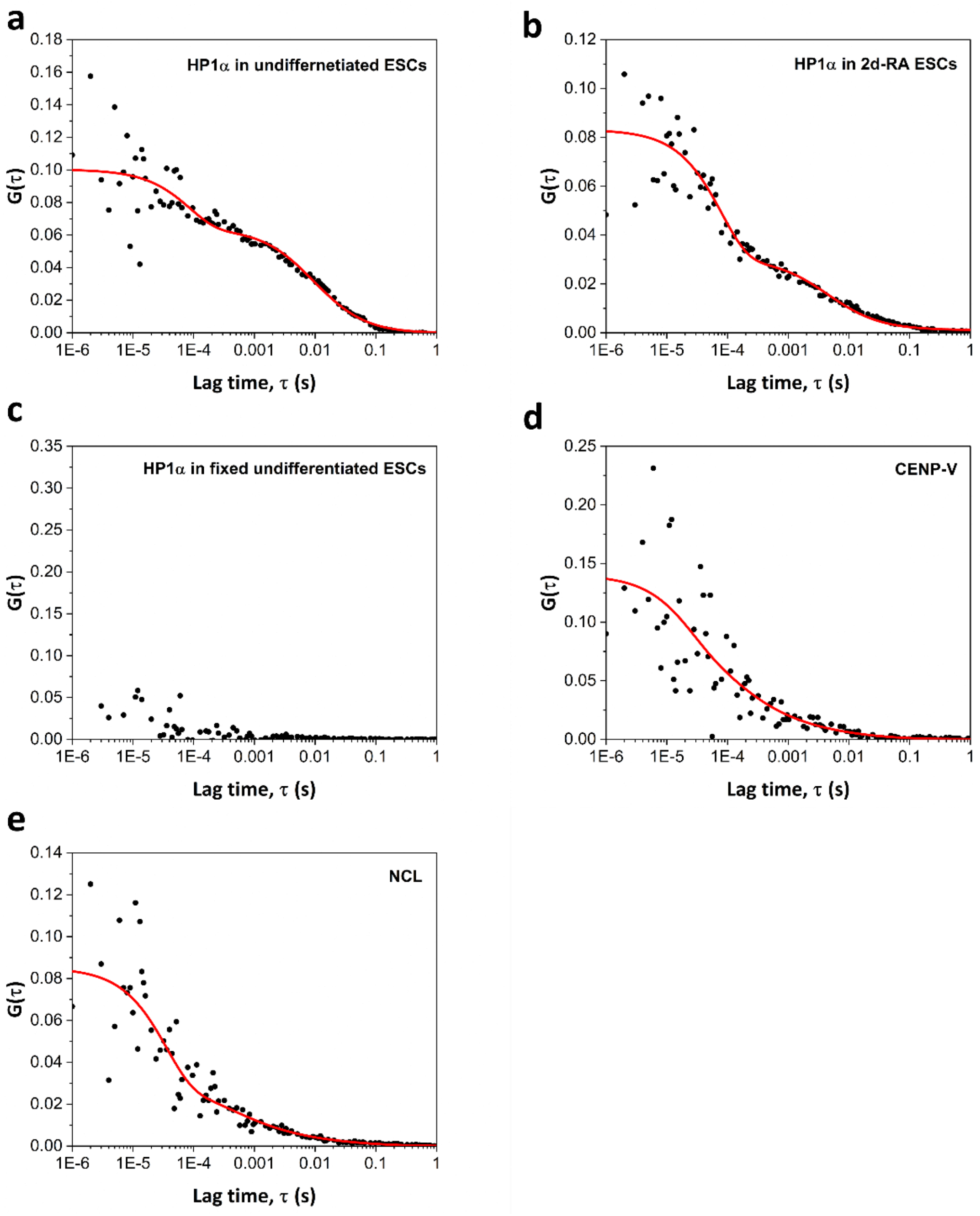


**Supplementary figure 3. Fluorescence correlation spectroscopy of mCherry-HP1α, -CENPV and -NCL inside biocondensates.** The fluorescence correlation curves of the different clusters measured inside their relevant biocondensates mCherry-HP1α in undifferentiated (a), 2d-RA (b) and fixed undifferentiated ESCs (c). mCherry-CENP-V (d) and mCherry-NCL in ESCs (e). The fluorescence correlation curves were recovered after reaching the steady-state fluorescence window during continuous photobleaching. Black dots are experimental data while the red lines are theoretical fit (see Supplementary table 2).

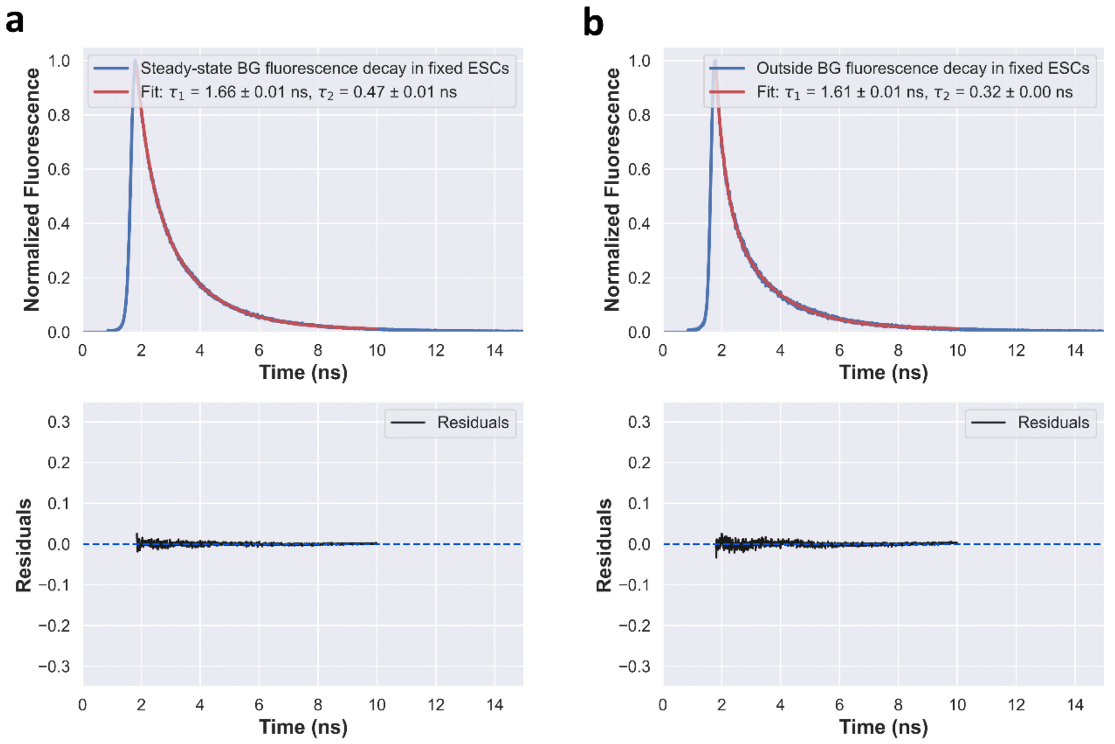


**Supplementary figure 4. Fitting results to fluorescence decays of mCherry-HP1α in fixed undifferentiated mouse ESCs.** The fluorescence decay curves and the fitting results, including residuals, of mCherry-HP1α from inside (a) and outside (b) heterochromatin biocondensates after reaching steady-state background.

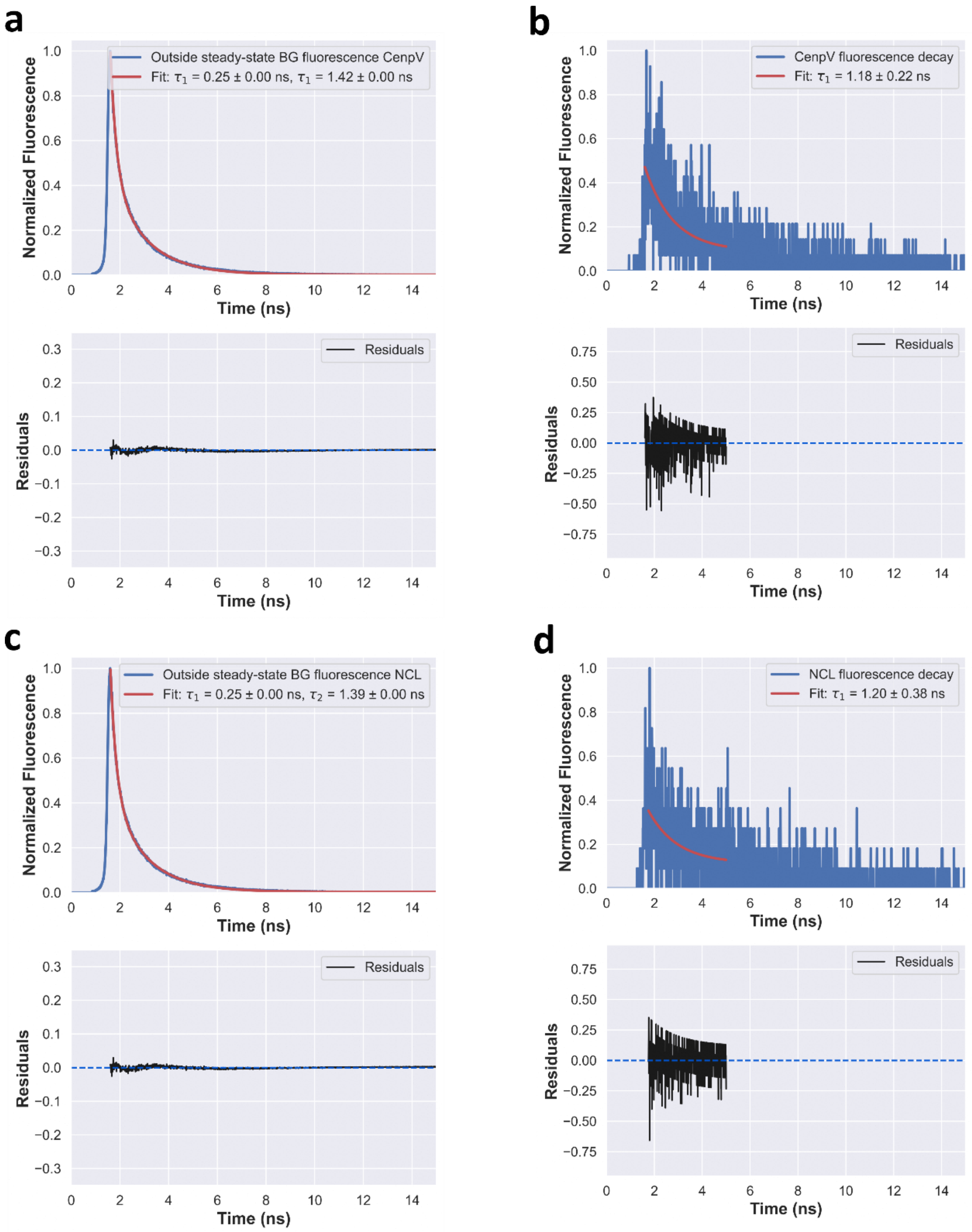


**Supplementary figure 5. Fitting results to fluorescence decays of mCherry-CENPV and mCherry-NCL.** The fluorescence decay curves of mCherry-CENPV (a) and mCherry-NCL (b) and the fitting results, including residuals, of fluorescence recovered from outside biocondensates. The fluorescence decay curves and fitting results, including residuals, of recovered mCherry-CENPV (c) or mCherry-NCL (d) fluorescent bursts from inside of their relevant biocondensates.

### Supplementary tables

#### Supplementary table 1: PyBroMo simulations of conditions similar to reaching steady-state conditions after continuous photobleaching to test in which conditions clusters can be identified as bursts above a background of diffuse molecules.

| Diffuse molecules | | | Cluster molecules | | | OBG[4] (kHz) | F=4 | | F=6 | | F=11 | | F=21 | |
|---|---|---|---|---|---|---|---|---|---|---|---|---|---|---|
| C[1] (nM) | D[2] (μm²/s) | B[3] (kHz) | C (pM) | D (μm²/s) | B (kHz) | | # Bursts | % clusters | # Bursts | % clusters | # Bursts | % clusters | # Bursts | % clusters |
| 3.1 | 100 | 30 | 62 | 1 | 300 | 2.5-3.0 | 942 | 60 | 776 | 76 | 767 | 86 | 877 | 92 |
| | | | | | 3,000 | | 3,726 | 76 | 3,514 | 83 | 3,825 | 91 | 3,645 | 93 |
| 0.5 | | | 52 | | 300 | 1.0-1.2 | 3,798 | 16 | 2,543 | 21 | 1,250 | 31 | 722 | 52 |

[1] Concentration
[2] Diffusion coefficient
[3] Brightness
[4] Observed background rates

#### Supplementary table 2: Best-fit models of FCS data using one-component diffusion models with a dynamic photophysical process.

| | D $\left(\frac{\mu m^2}{s}\right)$ [1] | $\tau_D (ms)$ | N [2] | $T$ [3] | $\tau_T$ (s)[4] |
|---|---|---|---|---|---|
| **mCherry-HP1α** in undifferentiated ESCs | $2.35 \pm 0.86$ [5] | $9.57 \pm 3.50$ | $9.97 \pm 0.50$ [6] | $0.36 \pm 0.04$ [7] | $8.5E^{-5} \pm 2.6E^{-5}$ [8] |
| **mCherry-HP1α** in 2d-RA ESCs | $5.00 \pm 1.99$ | $4.50 \pm 1.79$ | $12.18 \pm 757.32$ | $0.64 \pm 39.85$ | $8.0E^{-5} \pm 2.5E^{-4}$ |
| **mCherry-CENPV** | $0.29 \pm 3.18$ | $77.59 \pm 850.81$ | $7.15 \pm 0.78$ | $0.25 \pm 0.65$ | $2.4E^{-5} \pm 3.7E^{-5}$ |
| **mCherry-NCL** | $0.08 \pm 1.80$ | $281.25 \pm 6328.12$ | $11.74 \pm 0.91$ | $0.65 \pm 0.13$ | $3.7E^{-5} \pm 1.1E^{-5}$ |

[1]The diffusion coefficient calculated using eq. 5 (see Methods) after fitting the autocorrelation function
[2]The diffusion times are calculated using eq. 7 (see Methods)
[2]The number of fluorescent molecules inside the effective excitation volume eq. 5 (see Methods)
[3]The equilibrium fraction of the dynamic photophysical process eq. 5 (see Methods)
[4]The relaxation time of the dynamic photophysical process eq. 5 (see Methods)
[5,6,7,8]Fitting standard errors of the different parameters
